# Paradoxical effects of cigarette smoke and COPD on SARS-CoV2 infection and disease

**DOI:** 10.1101/2020.12.07.413252

**Authors:** M. Tomchaney, M. Contoli, J. Mayo, S. Baraldo, L. Shuaizhi, CR. Cabel, DA. Bull, S. Lick, J. Malo, S. Knoper, SS. Kim, J. Tram, J. Rojas-Quintero, M. Kraft, J. Ledford, Y. Tesfaigzi, F.D. Martinez, C. A. Thorne, F. Kheradmand, S. K. Campos, A. Papi, F. Polverino

## Abstract

**Introduction:** How cigarette smoke (CS) and chronic obstructive pulmonary disease (COPD) affect severe acute respiratory syndrome coronavirus 2 (SARS-CoV-2) infection and severity is controversial. We investigated the protein and mRNA expression of SARS-CoV-2 entry receptor ACE2 and proteinase TMPRSS2 in lungs from COPD patients and controls, and lung tissue from mice exposed acutely and chronically to CS. Also, we investigated the effects of CS exposure on SARS-CoV-2 infection in human bronchial epithelial cells.

**Methods:** In Cohort 1, ACE2-positive cells were quantified by immunostaining in FFPE sections from both central and peripheral airways. In Cohort 2, we quantified pulmonary ACE2 protein levels by immunostaining and ELISA, and both ACE2 and TMPRSS2 mRNA levels by RT-qPCR. In C57BL/6 WT mice exposed to air or CS for up to 6 months, pulmonary ACE2 protein levels were quantified by triple immunofluorescence staining and ELISA. The effects of CS exposure on SARS-CoV-2 infection were evaluated after 72hr in vitro infection of Calu-3 cells. After SARS-CoV-2 infection, the cells were fixed for IF staining with dsRNA-specific J2 monoclonal Ab, and cell lysates were harvested for WB of viral nucleocapsid (N) protein. Supernatants (SN) and cytoplasmic lysates were obtained to measure ACE2 levels by ELISA.

**Results:** In both human cohorts, ACE2 protein and mRNA levels were decreased in peripheral airways from COPD patients versus both smoker and NS controls, but similar in central airways. TMPRSS2 levels were similar across groups. Mice exposed to CS had decreased ACE2 protein levels in their bronchial and alveolar epithelia versus air-exposed mice exposed to 3 and 6 months of CS. In Calu3 cells in vitro, CS-treatment abrogated infection to levels below the limit of detection. Similar results were seen with WB for viral N protein, showing peak viral protein synthesis at 72hr.

**Conclusions:** ACE2 levels were decreased in both bronchial and alveolar epithelial cells from uninfected COPD patients versus controls, and from CS-exposed versus air-exposed mice. CS-pre-treatment did not affect ACE2 levels but potently inhibited SARS-CoV-2 replication in this in vitro model. These findings urge to further investigate the controversial effects of CS and COPD on SARS-CoV2 infection.

## Introduction

How cigarette smoke (CS) and chronic obstructive pulmonary disease (COPD) affect susceptibility and outcomes in SARS-CoV-2 infection is controversial. SARS-CoV-2 uses the angiotensin-converting enzyme 2 (ACE2) as the main host cell entry receptor^1^, and viral spike priming by proteases like TMPRSS2^2^. ACE2 expression is highest in the conducting airways, waning in the more distal bronchiolar and alveolar lung regions^3^. There have been mixed reports on the effect of CS and COPD on SARS-CoV-2 infection^4^, with most studies showing an up-regulation of ACE2 in clinical and experimental models of smoke-induced lung disease^5-10^. In this study, we evaluated: 1) ACE2 and TMPRSS2 expression in the airways of smokers with COPD, compared to smoker and never smoker (NS) controls, and 2) ACE2 expression in lungs of mice acutely and chronically exposed to CS; and 3) the effect of CS on SARS-CoV2 infection of bronchial epithelial cells *in vitro*.

## Methods

Lung tissue from two separate COPD cohorts and controls were studied to validate the results. Cohort 1: smokers with pulmonary nodules who had lung resection at Ferrara University Hospital in Italy; Cohort 2: patients who underwent surgery for lung nodules or lung transplant at the University of Arizona. Explanted lungs from NS who died of extrapulmonary causes from the Arizona Donor Network (both from Tucson, AZ) were used as control lungs.

In Cohort 1, ACE2-positive cells were quantified by immunostaining in FFPE sections from both central and peripheral airways. In Cohort 2, we quantified pulmonary ACE2 protein levels by immunostaining and ELISA, and both ACE2 and TMPRSS2 mRNA levels by RT-qPCR. In C57BL/6 WT mice exposed to air or CS for up to 6 months^11^ (Brigham and Women’s Hospital, IACUC-approved), pulmonary ACE2 protein was quantified by immunostaining and ELISA.

The effects of CS exposure on SARS-CoV-2 infection were evaluated after 72hr *in vitro* infection of Calu-3 cells^12^. Cells were sham- or CS extract-treated for 24hr before infection with SARS-CoV-2 isolate (USA-WA1/2020). Cells were either fixed for IF staining, or lysates/supernatants were harvested for WB/ELISA of the viral nucleocapsid (N) protein and ACE2. IF staining for dsRNA intermediates which arise during viral RNA (vRNA) replication, was performed with dsRNA-specific J2 mAb^13^, and infection was quantified by automated microscopy using the Nikon Eclipse Ti2 system.

The statistical methods are listed in the figure legend.

## Results

The average age of the Cohort 1 was 71±7 years; 33% were females and 21% were current smokers. COPD patients (N=10) had lower FEV1% compared to NS (N=7) and smokers (N=16) subjects (66±10 vs 99±10 and vs 104±18% predicted, respectively; p<0.001) and lower FEV1/FVC ratio (0.61±0.08 vs 0.78±0.07 and vs 0.81±0.10, respectively; p<0.001). The average age of the Cohort 2 was 66±9 years; 44% were females and 23% were current smokers. Never smokers (N=10) were younger (56±18 years; p<0.01) compared to smokers (N=16) and COPD patients (N=21) (67±6 and 66±9 years, respectively). COPD patients had lower FEV1% compared to NS and smokers (40±24 vs 77±16and vs 88±21% predicted, respectively; p<0.001) and lower FEV1/FVC ratio (0.41±0.18 vs 0.82±0.07 and vs 0.80±0.09, respectively; p<0.001). No difference was found in pack/years and smoking history (current vs former) between COPD and smokers without COPD in both cohorts. All the former smokers had quit smoking at least one year before the study.

In Cohort 1, a greater ACE2 expression was present in central vs. peripheral airways, but no difference was found in ACE2 expression in COPD central airways (% ACE2+ cells/bronchial epithelial cells [mean ± SEM]= 31%±11) vs. smoker and NS controls (22%±6 and 39%±16, and *P*=0.6 and 0.9, respectively, **Fig. 1A**). In contrast, COPD patients had a lower ACE2 expression (5%±2) in peripheral airways vs. both smoker and NS controls (19%±6; 21%±6, and *P* = 0.04 and P = 0.01, respectively). Similarly, in Cohort 2 we found both alveolar (mainly type II cells, ATII, **Fig. 1B** and **D**) and bronchiolar (mainly club cells and goblet cells, **Fig. 1C** and **D**) ACE2 protein expression levels, and ACE2 mRNA pulmonary levels (**Fig. 1E**) to be lower in COPD patients vs. both smoker and NS controls. We did not observe any difference between smoking status (current vs former smokers) with or without COPD (**Fig. 1E**, red circles), nor between COPD GOLD stages (not shown). After adjusting for age and sex, ACE2 mRNA levels were all still significantly lower (*P* < 0.05) in COPD lung vs. smoker and NS controls. The ACE2 mRNA expression significantly correlated with its protein measured by both IF and ELISA across all the analyzed subjects. At variance with ACE2, TMPRSS2 mRNA were not different among groups (*P* = 0.9 for all comparisons).

**Figure 1:**
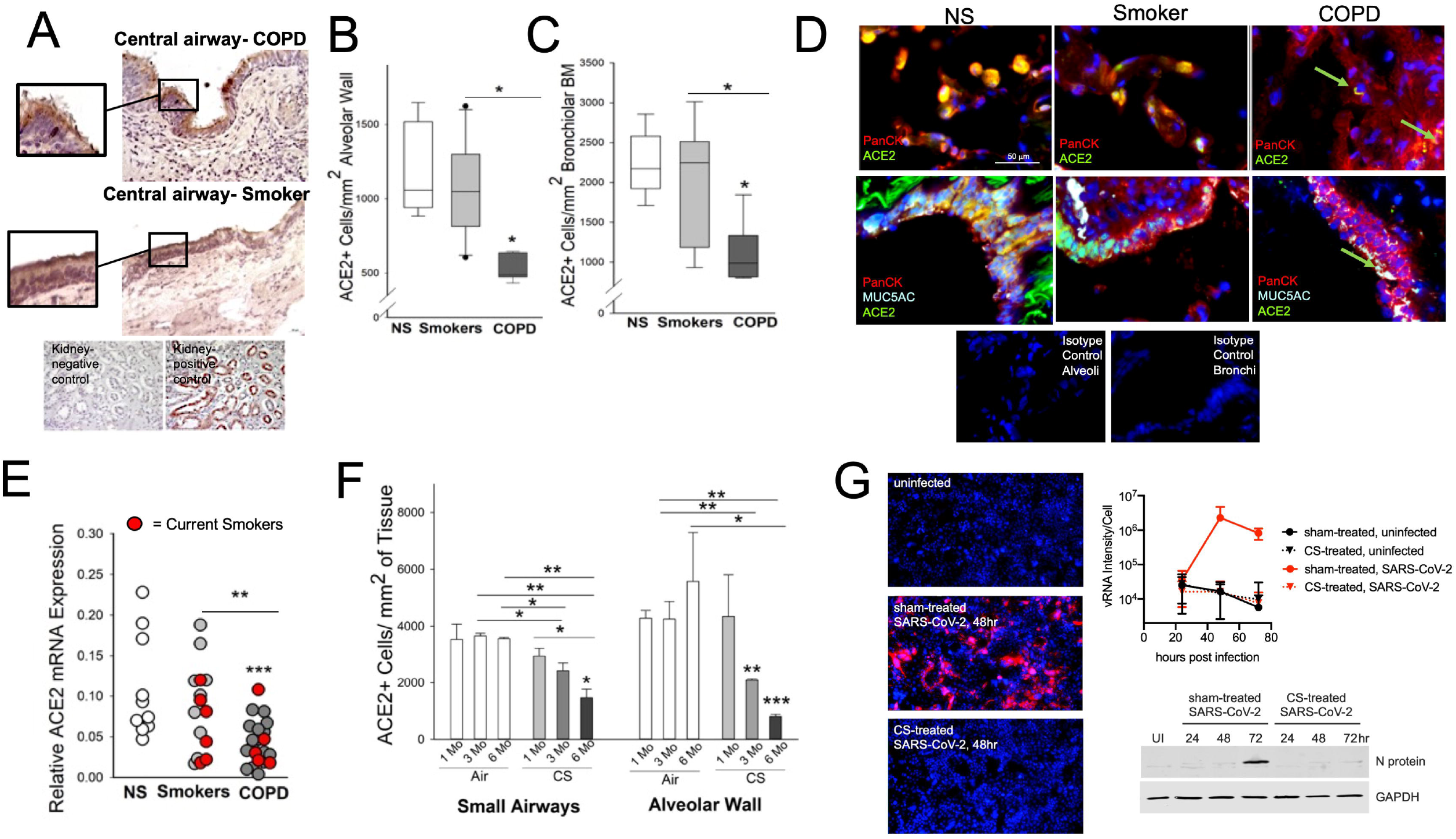
ACE2 expression in bronchial and alveolar epithelium from COPD patients, smoker and never-smoker (NS) controls and CS-exposed mice, and *in vitro* findings. The number of ACE2+ cells in the central airway bronchial epithelium was similar between patients with chronic obstructive pulmonary disease (COPD), smokers, and NS controls. In **A**, representative IHC for ACE2 images of central airways of a COPD patient (upper panel) and a smoker without COPD. The insets show details of the ciliated bronchial epithelium. The number of ACE2+ cells in the alveolar epithelium (**B**) and peripheral airway epithelium (**C**), normalized for length of the alveolar wall or basement membrane, respectively, was lower in patients with COPD versus smokers and NS controls. In **D**, triple immunofluorescence representative images of alveolar (upper panels) and bronchiolar epithelium (lower panels) from a COPD patient, a smoker without COPD, and a never-smoker (NS) where ACE2 staining is identified by green fluorochrome, the epithelium is identified by red fluorochrome, and the color yellow is obtained by merging the two fluorochromes. In **E**, the levels of ACE2 mRNA from peripheral lung samples was decreased between patients with chronic obstructive pulmonary disease (COPD) versus both smoker and NS controls. The red circles indicate the current smokers among the smoker controls and COPD patients. In **F**, WT C57BL6 mice were exposed to air or CS (CS) for up to 6 months (n=3/4 group); the number of ACE2+ cells in the bronchial epithelium was decreased in 6-month CS-exposed mice versus 1-, 3-, and 6-month air-exposed mice. The number of ACE2+ cells in the bronchial epithelium was decreased in 3-month CS-exposed mice compared to 3- and 6-month air-exposed mice. The number of ACE2+ cells in the bronchial epithelium was decreased in 6-month CS-exposed mice compared to 1-month CS-exposed mice. In **G**, we performed a 72hr in vitro infection of Calu-3 lung adenocarcinoma cells (n = 3 separate replicate experiments). Cells were sham- or CS-treated for 24hr prior to SARS-CoV-2 infection (2h viral infection in normal media, then remove inoculum). At indicated times, cells were either paraformaldehyde fixed for IF staining with DAPI and dsRNA-specific J2 antibody, or cell lysates were collected for SDS-PAGE. Automated microscopy with the Nikon Eclipse Ti2 system was used to quantify average J2 intensity per cell (DAPI stained nuclei), over the course of infection. Representative micrographs show infections at 48h, and western blot with shows viral N protein levels during the infection. Bar graphs and line graphs show means, scatter plots show every single value in the study population, boxes in box plots show the median values at 25^th^ and 75^th^ percentiles, and error bars show the 10^th^ and 90^th^ percentiles. * = P < 0.05; ** = P < 0.01; *** = P < 0.001 vs. NS (B, C, and E) or 1 month CS-exposed mice (F) or the group indicated. One-way analysis of variance tests was used for continuous variables and Z tests or Chi-square tests for categorical variables. For pairwise comparisons, parametric and nonparametric data were analyzed using two-sided Student’s t tests and Mann-Whitney U tests, respectively. Correlation coefficients were calculated using the Pearson or the Spearman rank method or the Dubin-Watson statistical correlation test for nonlinear data. P less than 0.05 was considered statistically significant. Analyses were performed using SPSS Statistical software and GraphPad Prism.

In mice, we found that the longer the CS exposure, the lower the levels of ACE2 staining in both bronchial and alveolar epithelial cells (**Fig. 1F**). As expected, the mice exposed to CS for 6 months had developed a COPD-like phenotype with airspace enlargement and small airway remodeling^11^, and had significantly higher MUC5AC+ bronchial cell numbers vs. air-exposed mice. Similarly, CS-exposed mice had a lower % ACE2+ MUC5AC+/ total MUC5AC+ cells in the bronchial epithelium vs. air-exposed mice (*P* < 0.05 for all comparisons, not shown). ELISA-measured pulmonary ACE-2 protein levels were significantly lower in 1-month CS-vs. air-exposed mice (*P* < 0.001, not shown).

*In vitro* SARS-CoV-2 infection of Calu3 cells was readily observed by dsRNA staining (**Fig 1G**). Whereas replication of vRNA peaked at 48hr in sham-treated cells, CS-treatment abrogated infection to levels below the detection limit. Similar results were seen with viral N protein expression, showing peak viral protein synthesis at 72hr (**Fig. 1G**). ACE2 protein levels were undetectable in the supernatants, whereas they were unchanged in CS-treated vs. sham cell lysates (not shown).

## Conclusion

The effects of CS and COPD on SARS-CoV-2 infection and disease a still a matter of debate. Herein we show that CS and COPD are associated with lower ACE2 levels in both human and murine cohorts, and CS-pre-treatment potently abrogates SARS-CoV-2 replication in this *in vitro* model. Our findings in human samples are not aligned with some data recently-published ^5-9^. Zhang et al.^6^ assessed the ACE2 gene expression in NS and smokers. Consistently with theirs and others’ findings^5,6,10^, we found ACE2 expression to be higher in central airways. While Zhang et al. did not include COPD patients in their analyses, Cai et al.^5^ and Smith et al.^9^ analyzed several datasets of gene expression of COPD and control small and large airway epithelium samples, concluding that ever-smoking was the main factor associated with high ACE2 levels; however, the effect of COPD was not consistent across datasets. Leung et al.^7^ reported higher pulmonary ACE2 protein and mRNA levels in smokers vs. NS and COPD vs. non-COPD. While ACE2 mRNA levels were quantified in subsegmental airway brushings, we assessed peripheral lung resections. Leung et al. also showed a diffuse increased ACE2 staining in bronchiolar epithelial cells from COPD vs. controls, while we detected apical staining in bronchiolar cells, with diffuse expression only in goblet and club cells. Jacobs et al.^8^ reported higher ACE2 protein and mRNA levels from ever smokers without and with COPD vs. NS. However, there are several differences in the methods (e.g., primers for different ACE2 splice variants) and study population (higher prevalence of hypertension, treatment with oral corticosteroids, LABA and LAMA, all known to affect ACE2 levels^14^, and a much higher percentage of females and current smokers) employed in our study. Finally, soluble ACE2 can be released from the epithelial surface into the airway lumen via sheddase cleavage, and thus, a dynamic expression of ACE2 in the airways in response to noxious stimuli such as CS and COPD could underlie such a variability in the findings.

CS-exposed mice had decreased ACE2+ cells in both alveolar and bronchial epithelial cells versus air-exposed mice, with the lowest number of ACE2+ cells found after six months of CS exposure. A previous study showed a lower ACE2 mRNA expression in the lungs of CS-exposed rats with a COPD-like phenotype compared with that in air-exposed animals^15^. In contrast, in a recent study, higher pulmonary ACE2 levels were found in CS-exposed vs. air-exposed mice^9^, but the smoke exposures were shorter than the ones used in our study, which might have resulted in less airway remodeling and emphysema. Further, cell subtypes expressing ACE2 in the CS-exposed lungs were not assessed.

Finally, we show for the first time that CS, while it does not change the ACE2 levels in bronchial epithelial cells, potently blocks SARS-CoV-2 replication in the same cells *in vitro*. Taken together, these studies, when conflated with features of attributable COVID-19 lung morbidity, indicate that lung ACE2 expression is a suboptimal marker of SARS-CoV-2 susceptibility and COVID-19 expression and morbidity, and additional factors likely play a role in the interaction between smoking, COPD, and SARS-CoV2 infection. We acknowledge that a major limitation of this study is the lack of insight into the mechanisms by which CS mediates SARS-CoV2 infection and disease, which we are currently investigating. Also, our pilot *in vitro* findings with acute CS exposure does not recapitulate our *in vivo* observations of decreased ACE2 levels in response to chronic CS.

In conclusion, we report in two independent human cohorts, lower protein and mRNA levels of ACE2 in both alveolar and bronchiolar epithelium of COPD patients versus smoker and NS controls. Consistently, we report that ACE2 levels were reduced in mice exposed chronically to CS versus air-exposed mice. Last, we report that CS pre-treatment potently abrogated SARS-CoV-2 replication in this *in vitro* model. These results point at complex biological interactions between CS, COPD, ACE2, and SARS-CoV2 entry in the host cells that need to be further explored in future clinical and translational studies.

## Acknowledgements

We are grateful to Jen Uhrlaub, for assistance the in the University of Arizona BSL3 suites, and to John Sullivan for assistance with the data analyses.

## Notes

**Sources of support:** Supported by funds from the Asthma and Airway Disease Research Center (University of Arizona); FP is supported by grants from the NIH/NHLBI (HL149744), Flight Attendants Medical Research Institute (#YFAC141004) and NIH/NHLBI (HL149744), and is recipient of the Parker B. Francis Fellowship; SB is supported by University of Padova DOR funds. FP. SKC is supported by a UA RII COVID-19 seed grant (#002196) and a grant from NIGMS (1R01GM136853). CAT is supported by UA TRIF grant and NIDDK R00 DK103126. FDM is supported by grants from NIH/NHLBI (HL139054, HL091889, HL132523, HL130045, HL098112, HL056177), the NIH/NIEHS (ES006614), the NIH/NIAID (AI126614), and the NIH/ Office of Director (OD023282).

### Competing Interest Statement

M. Contoli has received personal fees from Chiesi, AstraZeneca, Boehringer-Ingelheim, Alk-Abello, GSK, Novartis, Zambon, and scientific grants from Chiesi and University of Ferrara, Italy.
A. Papi: Board membership, consultancy, payment for lectures, grants for research, travel expenses reimbursement from GSK, AZ, Boehringer Ingelheim, Chiesi Farmaceutici, TEVA< Mundipharma, Zambon, Novartis, Menarini, Sanofi, Roche, Edmondpharma, Fondazione Maugeri, Fondazione Chiesi.

